# NOTCH1 signaling establishes the medullary thymic epithelial cell progenitor pool during mouse fetal development

**DOI:** 10.1101/600817

**Authors:** Jie Li, Julie Gordon, Edward L. Y. Chen, Luying Wu, Juan Carlos Zúñiga-Pflücker, Nancy R. Manley

## Abstract

The cortical and medullary thymic epithelial cell (cTEC and mTEC) lineages are essential for inducing T cell lineage commitment, T cell positive selection and the establishment of self-tolerance, but the mechanisms controlling their fetal specification and differentiation are poorly understood. Here, we show that Notch signaling is required to specify and expand the mTEC lineage. *Notch1* is expressed by and active in TEC progenitors. Deletion of *Notch1* in TECs resulted in depletion of mTEC progenitors and dramatic reductions in mTECs during fetal stages, consistent with defects in mTEC specification and progenitor expansion. Conversely, forced Notch signaling in all TEC resulted in widespread expression of mTEC progenitor markers and profound defects in TEC differentiation. In addition, lineage-tracing analysis indicated that all mTECs have a history of receiving a Notch signal, consistent with Notch signaling occurring in mTEC progenitors. Interestingly, this lineage analysis also showed that cTECs are divided between Notch lineage-positive and lineage-negative populations, identifying a previously unknown complexity in the cTEC lineage.

---

Notch signaling is a highly conserved pathway that plays a major role in the regulation of embryonic development and controls processes such as cell fate specification, differentiation and proliferation^1^. Notch is a transmembrane receptor protein, of which there are four (NOTCH1-4) in mammals. Importantly, Notch ligands are also membrane-bound, ensuring that ligand-receptor interactions can only occur between adjacent cells. Binding of a ligand to the receptor triggers a proteolytic event that cleaves the intracellular domain of the receptor, allowing it to enter the nucleus and regulate the expression of downstream genes.

The thymus is the primary lymphoid organ required for T cell production. The functional component of the thymus is comprised of thymic epithelial cells (TECs), which form a unique three-dimensional network that can be broadly divided into an outer cortex and an inner medulla. T cell differentiation takes place primarily via interactions between differentiating T cells and TECs, and a complete, organized and fully functional TEC compartment is essential for production of a diverse and self-tolerant T cell repertoire. Positive selection of T cells takes place in the cortex, where thymocytes capable of recognizing self-major histocompatibility complex (MHC) molecules are selected. The cells then enter the medulla and undergo negative selection to generate self-tolerant T cells that leave the thymus and enter the periphery. Notch signaling within lymphoid progenitor cells upon entry into the thymus is required for establishing T cell fate. Lymphocyte progenitors receive a NOTCH signal immediately upon entering the thymus, via interactions with the Delta-like 4 (Dll4) ligand on TECs^2^, that instructs them to commit to the T cell rather than alternative lineages^3^. NOTCH signaling is also required at multiple stages during T cell development for a variety of functions, including CD4 versus CD8 lineage commitment^4^. In addition to these critical and well-established roles in T cell differentiation, functional evidence has begun to emerge that suggests a role for NOTCH signaling in TECs. In addition to NOTCH ligands, TECs also express NOTCH receptors and pathway components^5,6^. Gain of function experiments suggest that NOTCH signaling is required to induce TEC development, particularly in the medullary lineage ^6,7^. These initial studies suggest that NOTCH signaling could play important roles in the differentiation of both the lymphoid and epithelial compartments. However, definitive *in vivo* experiments to establish the normal roles of NOTCH signaling in TEC development have not been performed.

All TECs have a single embryonic origin in the 3^rd^ pharyngeal pouch endoderm^8^, and functional studies suggest that TECs arise from a common thymic epithelial progenitor cell (TEPC)^9-11^. The precise developmental origin of TEC subsets is the subject of ongoing debate. There is evidence for both bipotent progenitors in the fetal mouse thymus^11,12^, and for lineage-specific progenitors for cortical TECs (cTECs)^13,14^ and medullary TECs (mTECs)^15,16^. There is also compelling evidence to suggest that a common progenitor population gives rise to mTEC lineage-specific progenitors^15^. Identifying key molecules involved in specification and maintenance of these different types of TEPCs will help to further elucidate how and when each lineage is specified during embryonic development.

We performed a series of loss-and gain-of-function and lineage tracing experiments to investigate the specific role of NOTCH1 signaling in fetal TEC development. Our results indicate that while all mTEC experience NOTCH signaling, only a subset of cTECs experience active NOTCH signaling, identifying a previously unappreciated aspect of cTEC differentiation. We also provide evidence of a requirement for NOTCH signaling in the establishment and maintenance/expansion of the mTEC progenitor pool in the fetal thymus.

## Results

### NOTCH1 activity in TEC progenitors in the fetal thymus

The NOTCH receptors and their downstream targets are expressed on TECs during late fetal development^6^ (see accompanying Liu, et al. paper). We first used immunohistochemistry (IHC) to assess NOTCH1 expression and activity, indicated by nuclear localization of cleaved NOTCH1, in the developing thymus. We first detected NOTCH1 in the nucleus in a few cells in the thymus primordium at E11.25, some of which were FOXN1^+^, and therefore TECs (Fig. 1A; white arrows). Thus, active NOTCH1 signaling was first detected in a few TECs around the time of initial *Foxn1* expression (E11.25), and is present in a subset of TECs at later stages. More FOXN1^+^ cells undergoing active NOTCH1 signaling were detected in the primordium just a few hours later (Fig. 1B), and were also present at E12.5 (Fig. 1C) and E14.5 (Fig. 1D). Next, we assessed Notch1 expression in TEC progenitors (TEPC) using an antibody against PLET1, a TEPC marker^9,10,17^. NOTCH1^+^FOXN1^+^PLET1^+^ TECs were detected in the thymus at E13.5 (Fig. 1E,F,G,H), suggesting that NOTCH1 signaling may play a role in early TEPCs during fetal thymus development.

**Figure 1.**
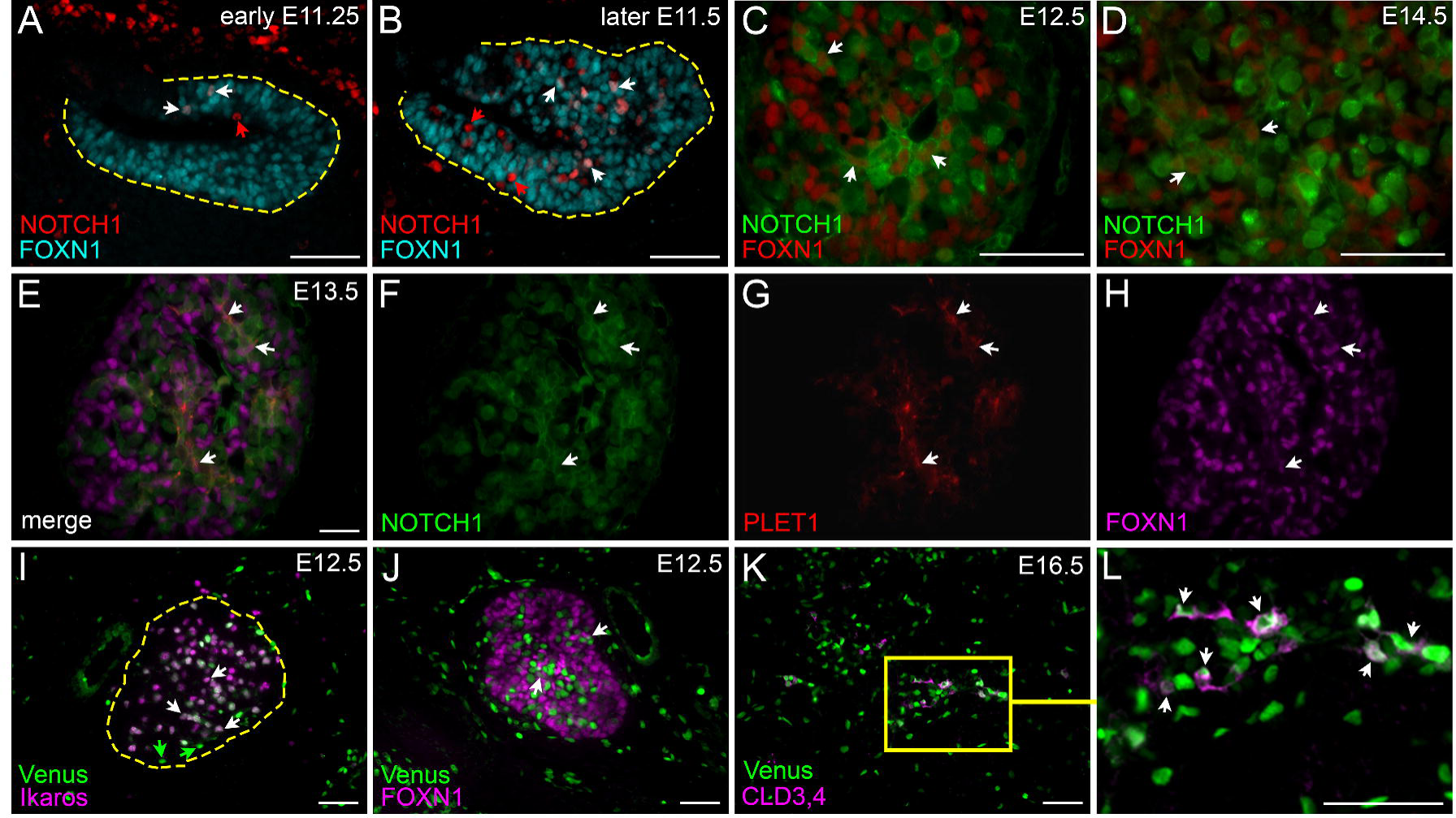
Notch1 expression and Notch activity in the fetal thymus. (A,B) Immunofluorescence of E11.25 (A) and E11.5 (B) wildtype thymus for cleaved NOTCH1 (red) and FOXN1 (cyan). White arrows in all panels indicate co-expressing cells; red arrows, NOTCH1^+^;FOXN1^-^ cells; dashed line outlines the primordium. (C,D) Immunofluorescence of E12.5 (C) and E14.5 (D) wildtype thymus for FOXN1 (red) and NOTCH1 (green). (E-H) Immunofluorescence of E13.5 wild type thymus for NOTCH1 (green), PLET1 (red), and FOXN1 (magenta). (I) Immunofluorescence of E12.5 CBF:H2B-Venus thymus for expression of IKAROS (magenta) and GFP (Venus; green). Green arrows, Venus expression in non-lymphocytes; dashed line outlines the thymus lobe. (J) Immunofluorescence of E12.5 CBF:H2B-Venus thymus for FOXN1 (magenta) and GFP (Venus; green). (K,L) Immunofluorescence of E16.5 CBF:H2B-Venus thymus for expression of Venus (green) and CLD3,4 (magenta). Box in (K) is zoomed area in (L). Scale bars, 50 μm. n > 3 for all experiments.

To further assess NOTCH signaling in the fetal thymus, we used a CBF:H2B-Venus transgenic mouse line^18^. These mice express nuclear localized Venus in cells undergoing active or recent NOTCH signaling. At E12.5, almost all of the Venus^+^ cells were Ikaros^+^ thymocytes undergoing active NOTCH signaling (Fig. 1I; white arrows), but a few Venus^+^Ikaros^-^ cells were also present at this stage (Fig. 1I; green arrows). Co-staining with FOXN1 confirmed that these were TECs (Fig. 1J). Claudin3,4 (CLD3,4) marks mTEC progenitors in the fetal thymus at mid-gestation^15^; at E16.5 nearly all CLD3,4^+^ cells expressed the CBF:H2B-Venus transgene (Fig.1K,L).

These data indicate that NOTCH1 signaling in TECs begins soon after the onset of *Foxn1* expression in a subset of cells that may represent progenitors, and that by E16.5 NOTCH1 signaling may act specifically in mTEPCs.

### Notch1 deletion in TEC results in fewer TEC progenitors in the fetal thymus

Since *Notch1* is expressed by a subset of fetal TECs, including potential mTEPCs, we used a loss-of-function approach to determine the role of NOTCH1 signaling in TEC differentiation. We used a *Notch1*^*flox*^ conditional allele^19^ together with a *Foxn1*^*Cre*^ deleter strain^20^ to remove NOTCH1 function from TECs at the onset of their differentiation.

To determine the effect of loss of *Notch1* on TEPC populations during fetal thymus development we performed IHC for PLET1^10^ and CLD3,4^15^. In control mice at E13.5, small clusters of PLET1^+^ cells were present in the thymus (Fig. 2A). In the *Foxn1*^*Cre*^*;Notch1*^*fx/fx*^ mutant thymus these clusters were rare and not always present (Fig. 2B). This phenotype was more severe at E16.5, when there were only a few PLET1^+^ cells in the mutant thymus (Fig. 2C,D). CLD3,4^+^ mTEC progenitors were also reduced at E13.5 and E16.5. Notably, in the control thymus, PLET1 and CLD3,4 were co-expressed at E13.5, whereas at E16.5 only a few cells co-expressed these markers (Fig. 2C; yellow arrows); most were positive for PLET1 or CLD3,4, but not both. These cells were arranged such that individual PLET1^+^CLD3,4^+^ double positive cells were surrounded by PLET1^+^ or CLD3,4^+^ single positive cells (Fig. 2C), rather than in homotypic clusters. Conversely, in the *Foxn1*^*Cre*^*;Notch1*^*fx/fx*^ mutant thymus, not only were there fewer PLET1 or CLD3,4 positive cells overall at E16.5, but all positive cells continued to express both PLET1 and CLD3,4 (Fig. 2D). The reduction in both the percentage and number of CLD3^+^ cells was confirmed by flow cytometry at E17.5 (*P* < 0.05) (Fig. 2E-G).

**Figure 2.**
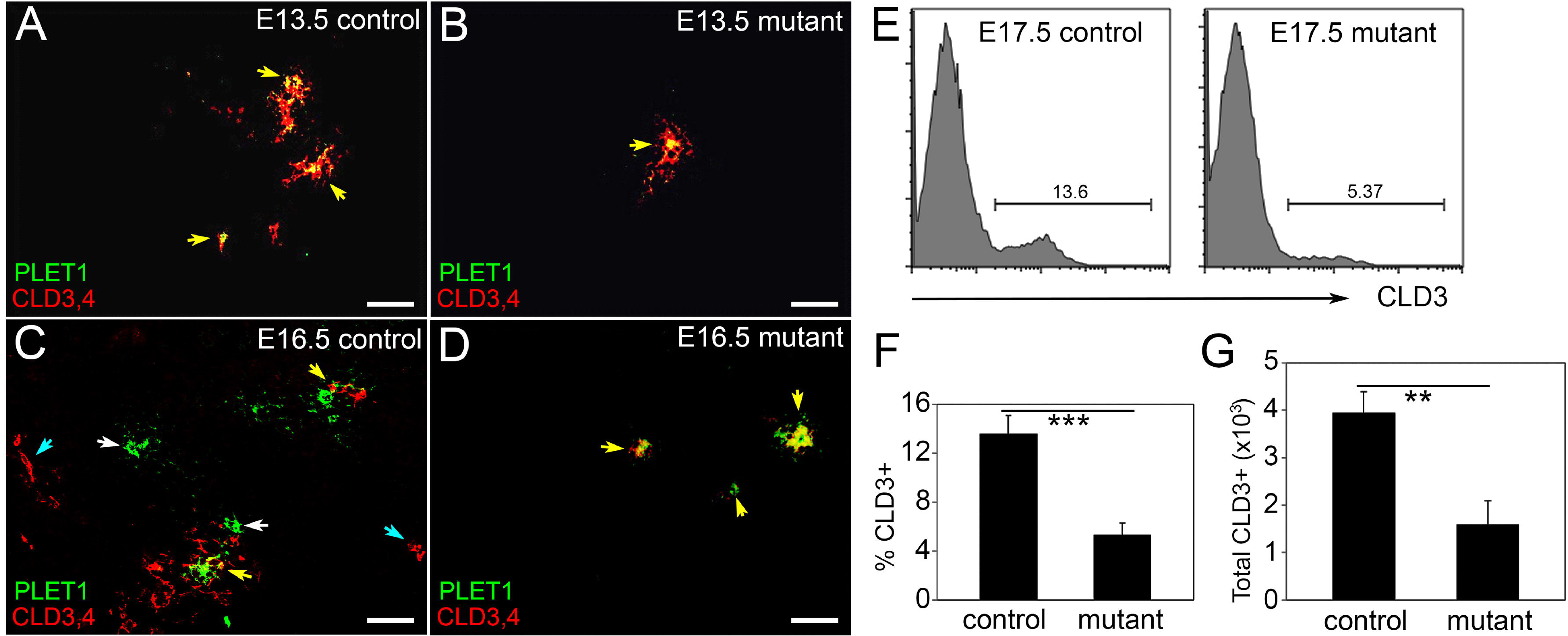
Notch1 deletion in TECs results in fewer TEPCs in the fetal thymus. (A-D) Immunofluorescence of E13.5 (A,B) and E16.5 (C,D) *Foxn1*^*Cre*^*;Notch1*^*fx/fx*^ mutant (B,D) and control (A,C) thymi for CLD3,4 (red) and PLET1 (green). White arrows, PLET1^+^;CLD3,4^-^ cells; cyan arrows, PLET1^-^;CLD3,4^+^ cells; yellow arrows, PLET1^+^;CLD3,4^+^ cells. (E) Histogram showing CLD3^+^ cells in *Foxn1*^*Cre*^*;Notch1*^*fx/fx*^ mutant and control thymi at E17.5. (E,F) Percentage and total number (G) of CLD3^+^ TECs in mutant and control thymi at E17.5. Scale bars, 50 μm. \*\*\**P* ≤ 0.001, \*\**P* ≤ 0.005. n > 3 for IHC; n = 5 for flow cytometry.

Together, these data show that *Notch1* deletion from TECs results in fewer putative fetal TEC progenitors, particularly mTEPCs, as shown by fewer PLET1 and CLD3,4 expressing cells. Furthermore, there were few or no PLET1^-^CLD3,4^+^ cells in the mutant thymus, suggesting a specific role for NOTCH1 in the lineage restriction of mTEC progenitors from a common progenitor during fetal thymus development.

### TEC differentiation and organization is abnormal in Foxn1^*Cre*^*;Notch1*^*fx/fx*^ *mutants*

To assess TEC differentiation and function after *Notch1* deletion, we performed IHC using a well-defined panel of markers that identify specific TEC subsets within the cortical and medullary compartments of the fetal thymus. We used Keratin 8 (K8), CD205 and β5t to label cTECs, and Keratin 5 (K5), Keratin 14 (K14), AIRE, and the lectin UEA1 to label mTEC subpopulations. In controls at E16.5, small distinct regions of K5, K14 and UEA1 positive cells mark the newly expanding medulla in the developing thymus (Fig. 3A,E,G). In the *Foxn1*^*Cre*^*;Notch1*^*fx/fx*^ mutant thymus, the medulla primarily consisted of one larger central region rather than several smaller islands (Fig. 3B,E,F,H). This phenotype was also seen at the newborn stage (not shown). Furthermore, there were dramatically fewer AIRE^+^ cells in the mutant thymus at E16.5 (Fig. 3C,D); an average of 21 cells per section for the control versus only one cell per section for the mutant, suggesting a nearly complete block in mTEC terminal differentiation. Flow cytometry confirmed the reduction in the number and frequency of mTECs in the *Foxn1*^*Cre*^*;Notch1*^*fx/fx*^ mutant thymus at E17.5 (*P* < 0.05; Fig. 3I).

**Figure 3.**
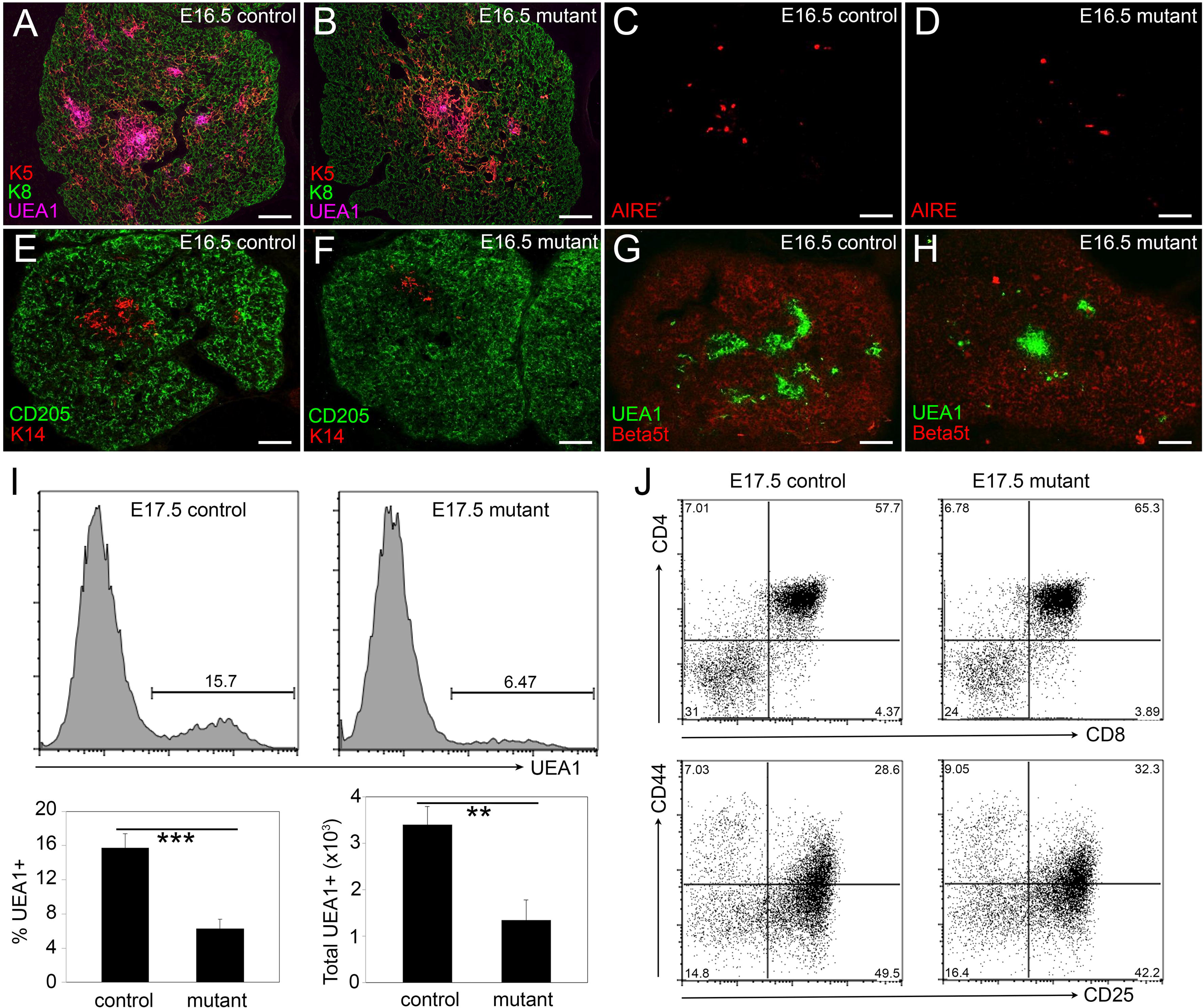
Notch1 deletion from TECs affects mTEC organization and differentiation. (A,B) Immunofluorescence of E16.5 *Foxn1*^*Cre*^*;Notch1*^*fx/fx*^ mutant (B) and control (A) thymus for K5 (red), K8 (green) and UEA1 (magenta). (C,D) Immunofluorescence of E16.5 *Foxn1*^*Cre*^*;Notch1*^*fx/fx*^ mutant (D) and control (C) thymus for AIRE. (E,F) Immunofluorescence of E16.5 *Foxn1*^*Cre*^*;Notch1*^*fx/fx*^ mutant (F) and control (E) thymus for K14 (red) and CD205 (green). (G,H) Immunofluorescence of E16.5 *Foxn1*^*Cre*^*;Notch1*^*fx/fx*^ mutant (H) and control (G) thymus for UEA1 (green) and β5t (red). (I) Flow cytometry showing histogram (top), percentage (bottom left) and total number (bottom right) of UEA1^+^ cells in *Foxn1*^*Cre*^*;Notch1*^*fx/fx*^ mutant and control thymi at E17.5. (J) Flow cytometric analysis of intrathymic thymocytes from E17.5 *Foxn1*^*Cre*^*;Notch1*^*fx/fx*^ mutant and control thymi stained for CD4, CD8, CD25 and CD44. Top panels show CD4 versus CD8; bottom panels show DN subsets with CD44 versus CD25. Scale bars, 50 μm. \*\*\**P* ≤ 0.001, \*\**P* ≤ 0.005. n > 3 for IHC; n > 5 for flow cytometry.

As total TEC numbers were similar in control and *Foxn1*^*Cre*^*;Notch1*^*fx/fx*^ mutants (*P* = 0.32), the reduction in mTEC frequency was correlated with a relative increase in cTECs. The relative cTEC frequency was significantly increased (controls, 84.26 +/-1.65; mutants, 93.71 +/-1.11; p= 0.0001), although cTEC numbers were not significantly different (*P* = 0.27). The cTEC markers β5t and CD205 expression appeared normal at E16.5 (Fig. 3E-H). Therefore, the primary defect in TEC based on this analysis was in the mTEC lineage.

As the TEC microenvironment governs thymocyte development, we determined whether the observed TEC defects affected thymocyte populations. Early T cell precursors express neither CD4 nor CD8, and are termed double-negative (DN) thymocytes. DN cells are subdivided into four differentiation stages (DN1, CD44^+^CD25^-^; DN2, CD44^+^CD25^+^; DN3, CD44^-^CD25^+^; and DN4, CD44^-^Cd25^-^). Interactions with cTECs and mTECs mediate positive and negative selection, generating CD4^+^ and CD8^+^ single-positive (SP) T cells. Intrathymic T cell development appeared normal in both *Foxn1*^*Cre*^*;Notch1*^*fx/fx*^ (Fig. 3J), as the percentages of these different subsets were not different between mutants and controls.

In summary, NOTCH1 deletion in TEC at the onset of *Foxn1* initiation affects TEC organization and mTEC differentiation, but does not obviously affect T cell development in the fetal thymus.

### Constitutive activation of Notch signaling in TECs leads to an increase in TEPCs and a block in mTEC differentiation

Given that *Notch1* deletion resulted in fewer TEPCs and an apparent block or reduction in mTEC differentiation, we predict that *Notch1* overexpression might have the opposite effect. We therefore activated NOTCH1 signaling in all TECs from the onset of their differentiation in gain-of-function experiments using a *Rosa*^*N1-IC*^ inducible strain^21^ activated by the *Foxn1*^*Cre*^ deleter strain^20^. In the *Rosa*^*N1-IC*^ mice, the NOTCH1 intracellular domain (N1-IC) targeted to the *Rosa26* locus; Cre-mediated deletion of a *loxp/stop/loxp* cassette results in heritable, constitutive expression of N1-IC, resulting in constitutive NOTCH1-mediated signaling. We analyzed *Foxn1*^*Cre*^*;Rosa*^*N1-IC*^ embryos using markers of TECs, TEPCs and developing T cells.

While K5 and K8 are markers for medullary and cortical TECs, respectively, cells that co-express these markers are thought to contain a progenitor population, and are normally located at the cortico-medullary junction^22^. In the control E14.5 thymus, proto-medullary areas were beginning to down regulate K8 in the center surrounded by a band of K8^+^K5^+^ cells, while the remainder of TEC were K5 negative, delineating the emerging cortical and medullary regions (Fig. 4A-C). However, in the *Foxn1*^*Cre*^*;Rosa*^*N1-IC*^ thymus at the same stage, almost all TECs were K8^+^K5^+^, with only a few single K8^+^ cells (Fig. 4D-F and inset). Furthermore, both PLET1 and CLD3,4 positive cells were expanded in the *Foxn1*^*Cre*^*;Rosa*^*N1-IC*^ thymus at E15.5 (Fig. 4G-N). Although PLET1 and CLD3,4 single positive cells were present in the *Foxn1*^*Cre*^*;Rosa*^*N1-IC*^ thymus, most of these cells expressed both markers. Flow cytometry at E15.5 showed about a 4-fold expansion in the frequency of CLD3^+^ cells in the *Foxn1*^*Cre*^*;Rosa*^*N1-IC*^ mutant thymus compared to littermate controls (*P* < 0.05; Fig. 4O), and the number of CLD3^+^ cells more than doubled in the mutant (an average of 1233 (SD = 387.1), versus 496 (SD = 50.1) cells in the controls (n = 3; *P* = 0.03). Total TEC cellularity was not different between mutant and control at this stage (*P* = 0.32). Flow cytometry for UEA1 also revealed a dramatic expansion of the medullary compartment in the *Foxn1*^*Cre*^*;Rosa*^*N1-IC*^ mutant thymus (Fig. 4P) (*P* < 0.05). This relative increase in progenitor-like phenotypes persisted at E18.5, by which time cysts lined with PLET1 and CLD3,4 positive cells had begun to appear (Fig. 4Q-X).

**Figure 4.**
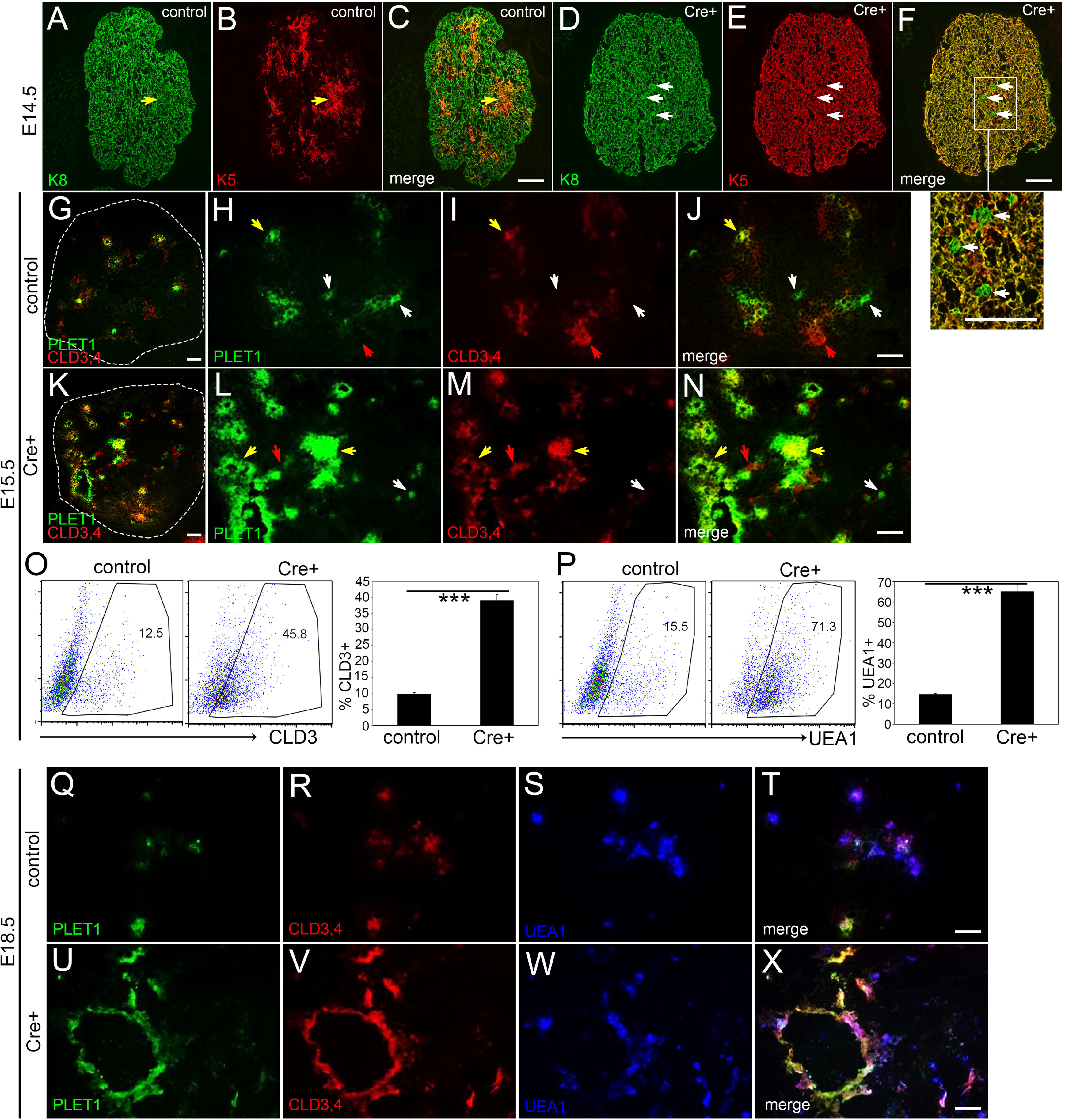
Notch1 activation in TECs causes an increase in the number of TEPCs at fetal stages. (A-F) Immunofluorescence of E14.5 *Foxn1*^*Cre*^*;Rosa*^*N1-IC*^ Cre^+^ (D-F) and control (A-C) thymus for K5 (red) and K8 (green). White arrows, K8^+^;K5^-^ cells; yellow arrows, K8^+^;K5^+^ cells. (G-N) Immunofluorescence of E15.5 control (G-J) and *Foxn1*^*Cre*^*;Rosa*^*N1-IC*^ Cre^+^ (K-N) thymus for PLET1 (green) and CLD3 (red). Dashed line in (G) and (K) outlines thymus lobe. White arrows, PLET1^+^;CLD3,4^-^ cells; red arrows, PLET1^-^;CLD3,4^+^ cells; yellow arrows, PLET1^+^;CLD3,4^+^ cells (O) Flow cytometric analysis of CLD3 expression in TECs from E15.5 *Foxn1*^*Cre*^*;Rosa*^*N1-IC*^ Cre^+^ and control thymi. (P) Flow cytometric analysis of UEA1 expression in TECs from E15.5 *Foxn1*^*Cre*^*;Rosa*^*N1-IC*^ Cre^+^ and control thymi. For (O, P), dot plots show one representative thymus; bar graph shows average values for 3 thymi. (Q-X) Immunofluorescence of E18.5 control (Q-T) and *Foxn1*^*Cre*^*;Rosa*^*N1-IC*^ Cre^+^ (U-X) thymus for PLET1 (green), CLD3 (red) and UEA1 (blue). Scale bars, 50μm. \*\*\**P* ≤ 0.001. n > 5 for IHC; n = 3 for flow cytometry.

mTEC differentiation did not occur normally in the *Foxn1*^*Cre*^*;Rosa*^*N1-IC*^ thymus. At E15.5, instead of the normal isolated islands of K14 expression (Fig. 5A), K14 was present throughout the mutant thymus, similar to K5 (Fig. 5B). There were also fewer and smaller clusters of UEA1^+^ cells (Fig. 5B,D) and very few AIRE^+^ cells compared to controls (Fig. 5C,D), indicating a block in mTEC terminal differentiation. By E18.5, this phenotype had progressed further. While widespread expression of K8, K5, and K14 showed that the thymus was still epithelial in nature, with (Fig. 5E-H), there was an almost complete absence of any recognizable organ structure at the newborn stage, as the epithelial network had essentially collapsed and the thymus was composed almost entirely of large cysts (Fig. 5I,J). Together, these data suggest that prolonged NOTCH1 signaling in TECs forces mTEC lineage commitment, but prevents differentiation, ultimately leading to a complete collapse of the TEC network.

**Figure 5.**
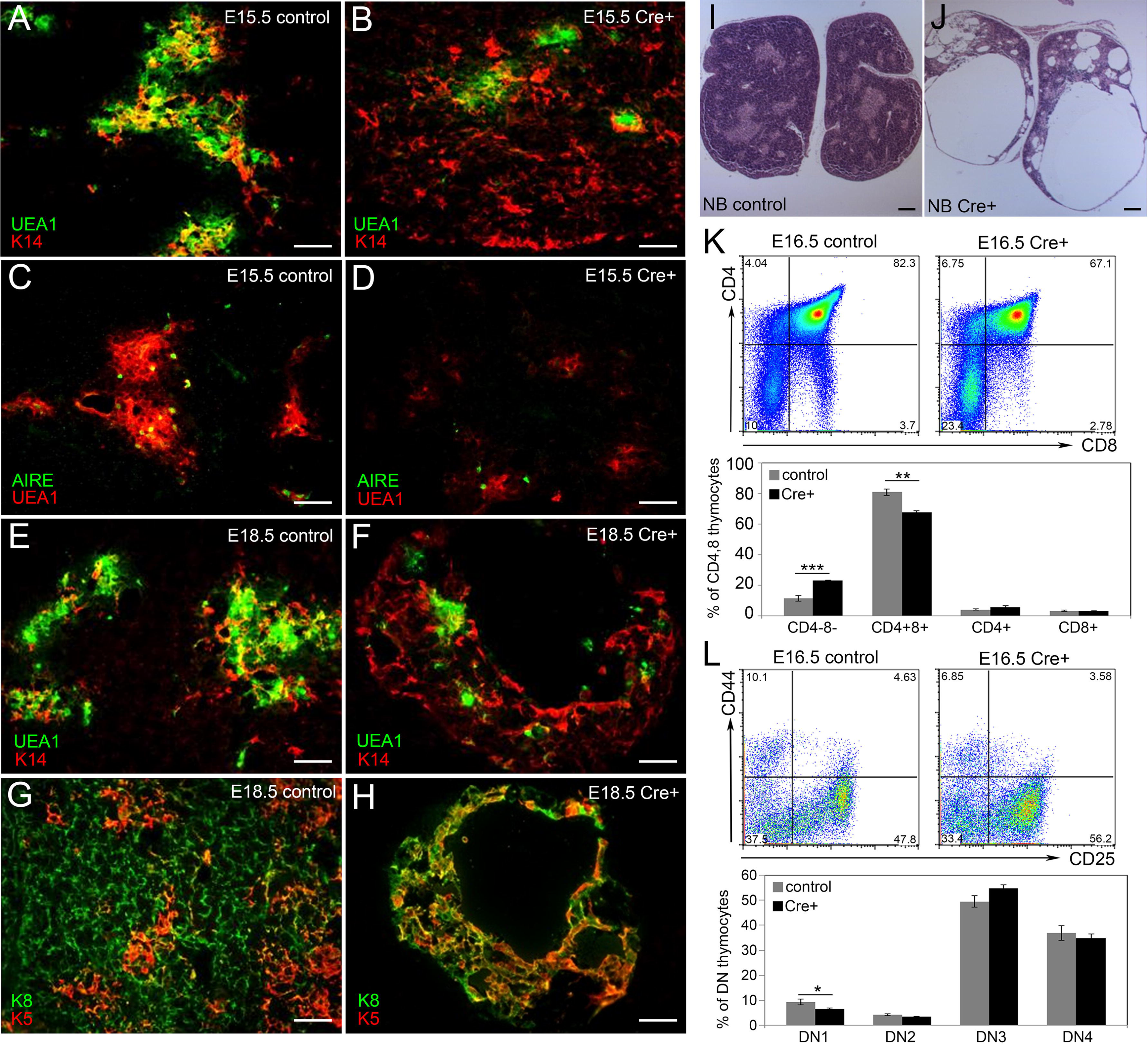
Ectopic expression of Notch1 in all TECs blocks fetal TEC differentiation and affects T cell development. (A,B) Immunofluorescence of E15.5 *Foxn1*^*Cre*^*;Rosa*^*N1-IC*^ Cre^+^ (B) and control (A) thymus for K14 (red) and UEA1 (green). (C,D) Immunofluorescence of E15.5 *Foxn1*^*Cre*^*;Rosa*^*N1-IC*^ Cre^+^ (D) and control (C) thymus for UEA1 (red) and AIRE (green). (E,F) Immunofluorescence of E18.5 *Foxn1*^*Cre*^*;Rosa*^*N1-IC*^ Cre^+^ (F) and control (E) thymus for K14 (red) and UEA1 (green). (G,H) Immunofluorescence of E18.5 *Foxn1*^*Cre*^*;Rosa*^*N1-IC*^ Cre^+^ (H) and control thymus for K5 (red) and K8 (green). (I,J) H&E staining of newborn (NB) *Foxn1*^*Cre*^*;Rosa*^*N1-IC*^ Cre^+^ (J) and control (I) thymus. (K) Flow cytometric analysis of thymocytes from E16.5 *Foxn1*^*Cre*^*;Rosa*^*N1-IC*^ Cre^+^ and control thymi stained for CD4 and CD8. (L) Flow cytometric analysis of thymocytes isolated from E16.5 *Foxn1*^*Cre*^*;Rosa*^*N1-IC*^ Cre^+^ and control thymi stained for DN subsets using CD44 and CD25. For (K,L), dot plots show one representative thymus for each genotype; bar graph shows average values for at least 5 thymi. Scale bars, 50 μm. \*\*\**P* ≤ 0.001, \*\**P* ≤ 0.005, \**P* ≤ 0.01. n > 5 for IHC; n > 5 for flow cytometry.

In contrast to the loss-of-function models, thymocyte development was affected by the abnormal TEC microenvironment in the *Foxn1*^*Cre*^*;Rosa*^*N1-IC*^ mice. The strongest effect was on total thymocyte numbers, which were reduced in the *Foxn1*^*Cre*^*;Rosa*^*N1-IC*^ thymus, with an average of 1.9×10^6^ thymocytes (SD = 0.51) in the mutant thymus compared with 12.5×10^6^ (SD = 2.12) in the control (*P* = 0.002). However, thymocyte differentiation was only mildly affected. Flow cytometry analysis of E16.5 *Foxn1*^*Cre*^*;Rosa*^*N1-IC*^ thymocytes revealed a slightly lower percentage of CD4^+^8^+^ cells (Fig. 5K), and an increase in DN3 (CD44^-^CD25^+^) cells (Fig. 5L) in the E16.5 mutant thymus compared to controls, suggesting a mild block at the DN3-DN4 transition. By late fetal stages, the thymic structure had deteriorated beyond the ability to support any thymocyte development.

Thus, dysregulation of NOTCH signaling throughout the TEC compartment during fetal development results in an abnormal TEC environment with an expanded mTEPC compartment, a major block to mTEC differentiation, and eventually causes complete collapse of the epithelial network. These data further support a role for NOTCH1 signaling in specifying the mTEPC pool during fetal development. These data also suggest that while NOTCH1 must be present for mTEPC specification, prolonged and/or excessive NOTCH1 signaling is detrimental to their differentiation.

### Mosaic deletion of Notch1 shows that mTEC specification requires NOTCH signaling

*Foxn1*^*Cre*^ initiates *Cre* expression at E11.25 ^20^, very similar to the timing with which mTEC specification may initiate ^23^, and coincident with our expression data showing that active NOTCH1 signaling in TECs in the developing thymus until E11.25 (Fig. 1A). Thus, it is possible that the few mTECs that are present in the *Foxn1*^*Cre*^*;Notch1*^*fx/fx*^ mutant thymus underwent specification prior to *Notch1* deletion. Since *Foxn1*^*Cre*^ is also active throughout TEC differentiation, these cells could have deleted *Notch* after mTEC specification; but since *Notch* expression is dispensable for or even detrimental to mTEC differentiation, this later deletion would have no effect. It would, however, make it impossible for us to determine whether this scenario was correct, as we cannot determine whether *Notch* was deleted before or after mTEC specification in these mTECs.

To test this possibility, we deleted *Notch1* from throughout the pharyngeal endoderm using *Foxa2*^*CreER*^ with a single pulse of tamoxifen at E8.5^24^, prior to the onset of *Foxn1* expression^25^. We have previously shown that this single pulse of CRE activity produces a mosaic deletion in the 3^rd^ pharyngeal pouch^26^, ideal for testing whether *Notch1* deleted cells can contribute to the mTEC lineage. *Foxa2*^*CreER*^*;Notch1*^*fx/fx*^ mice had fetal thymus phenotypes consistent with those obtained using *Foxn1*^*Cre*^, with reductions in both mTEC progenitor numbers and medullary size (Figs. S1, S2). Using PCR primers that selectively amplified either the undeleted or deleted allele, we performed qPCR on sorted cTEC and mTEC populations from Cre negative controls, *Foxa2*^*CreER*^*;Notch1*^*+/fx*^ heterozygotes, and *Foxa2*^*CreER*^*;Notch1*^*fx/fx*^ homozygous mutants (Fig. 6A-C). As expected for mosaic deletion, all cell populations from all genotypes were positive for the undeleted allele, and the band corresponding to the deleted allele was absent from Cre negative controls and present in all cell populations in heterozygotes (Fig. 6D). Strikingly, in *Foxa2*^*CreER*^*;Notch1*^*fx/fx*^ homozygous mutants only cTEC populations had the deleted allele, which was completely absent in mTECs (Fig. 6D). These data strongly support the conclusion that specification to the mTEC lineage requires NOTCH1 signaling, and is consistent with the idea that mTEC that are present in the *Foxn1*^*Cre*^*;Notch1*^*fx/fx*^ homozygous mutants had specified to the mTEC lineage prior to *Foxn1* expression.

**Figure 6.**
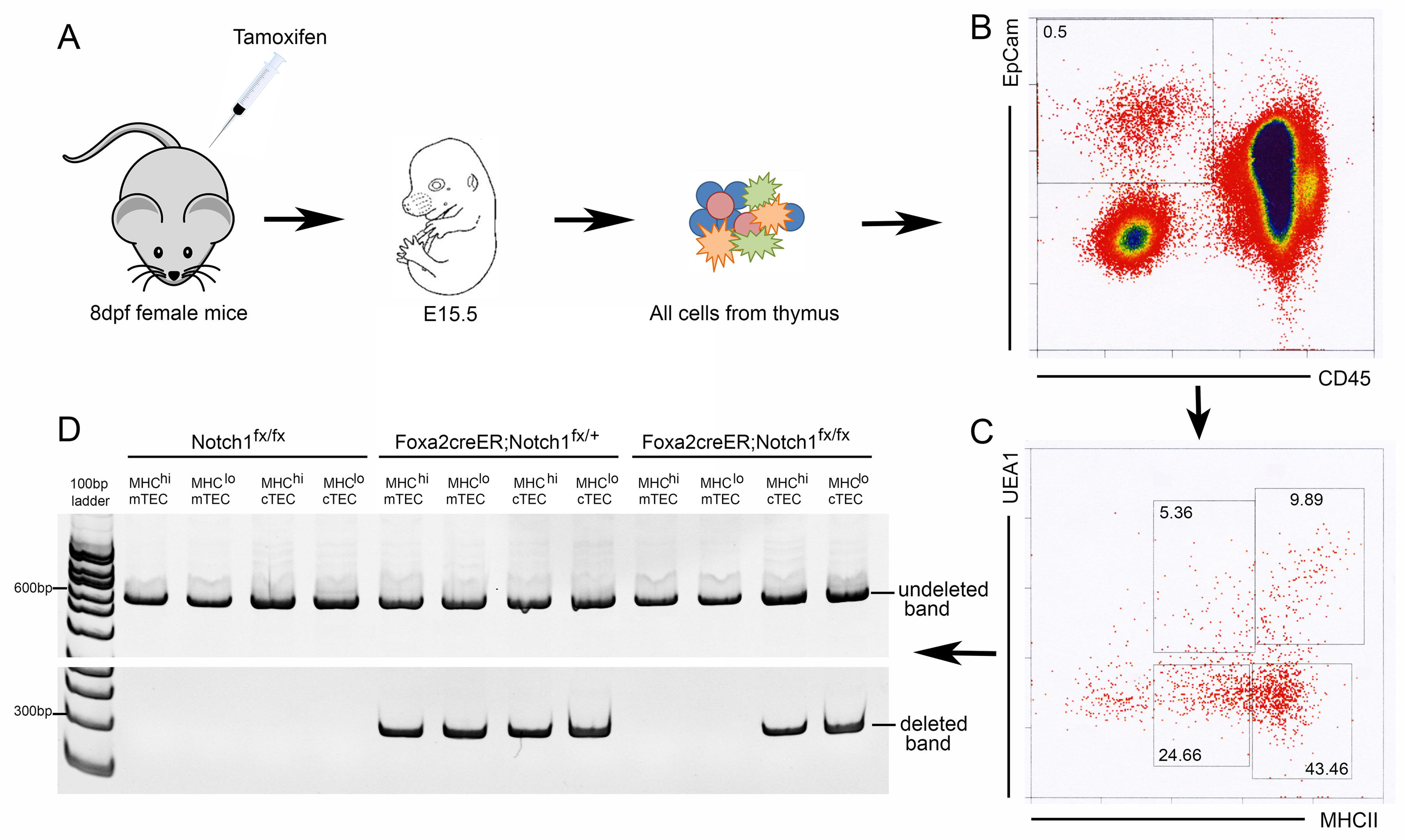
*Notch1-*deleted TEC are unable to contribute to the mTEC lineage. (A) Scheme for generating TECs with mosaic *Notch1* deletion for analysis. Pregnant dams are injected at E8.5 (8dpf), embryos are collected at E15.5, and the thymus dissected and dissociated into single cells. (B) Gating for isolation of EpCam+CD45-TECs. (C) Gating for MHCII^lo^ and MHCII^hi^ cTEC (UEA-1-) and mTEC (UEA-1+). (D) PCR of genomic DNA with primers specific for the wild-type undeleted and deleted alleles of *Notch1.* Genotypes and cell populations represented are indicated above each lane.

### Notch signaling is required in TECs at multiple fetal stages

The *Foxa2*^*CreER*^ and *Foxn1*^*Cre*^ experiments support previous data showing that mTEC begin to be specified quite early in thymus organogenesis, at around the time that *Foxn1* is first expressed, and that mTEC specification is *Foxn1*-independent^23^. To test the timing of *Notch1* requirement in TECs across fetal development, we utilized a genetic system in which the NOTCH pathway transcription factor RBPj is deleted in all TEC using *Foxn1*^*Cre*^, and then the capacity to respond to normal, physiological NOTCH signals is reactivated in a temporal and cell type specific manner using doxycycline-controlled expression of transgenic RBPj-HA (RBPj^fx/fx^;Foxn1^Cre^;Rosa^rtTA^;Tet^on^-RBPj-HA)^27^. *Rbpj* deletion using *Foxn1*^*Cre*^ resulted in similar phenotypes at E16.5 and NB stages as *Notch1* deletion, with many fewer mTEC, smaller medullary regions, and near complete loss of PLET1+ and CLD3,4+ cells (“un-induced”; Fig. 7B, E, H, and L panels) (see also companion paper, Liu et al.). We then temporally activated Notch signaling responsiveness in TEC by providing doxycycline from E0-E14 (assayed at E16 and NB), or from E14-NB (assayed at NB).

**Figure 7.**
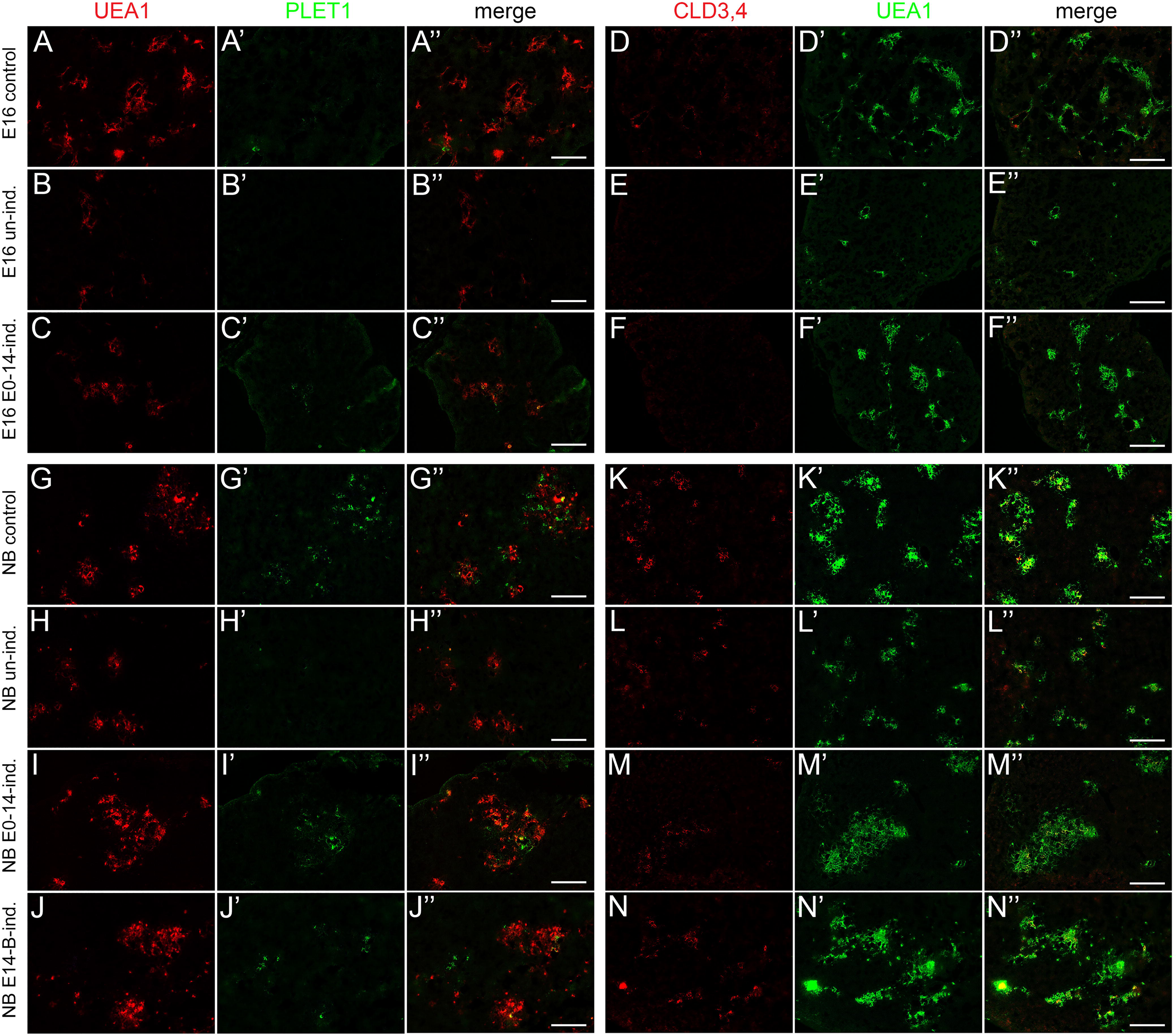
Analysis of the temporal requirement for NOTCH signaling in fetal TEC. Labels on the left refer to the entire row; marker names across the top refer to the entire column. In each row, panels with the same letter are single color or merged versions of the same image. A, A’, A” and D, D’, D”. Control RBPj^fx/+^;Foxn1^Cre^;Rosa^rtTA^;Tet^on^-RBPj-HA embryos collected at E16.5 have a wild-type phenotype. B, B’, B” and E, E’, E”. Uninduced RBPj^fx/fx^;Foxn1^Cre^;Rosa^rtTA^;Tet^on^-RBPj-HA (RBPj^ind^) embryos collected at E16 have a TEC-specific *Rbpj* null phenotype. C, C’, C” and F, F’, F”. RBPj^ind^ embryos injected with doxycycline daily from E0-E14 only, collected at E16. G, G’, G” and K, K’, K”. Control embryos collected at newborn (NB) stage. H, H’, H” and L, L’, L”. Uninduced RBPj^ind^ embryos collected at NB stage. I, I’, I” and M, M’, M”. RBPj^ind^ embryos injected with doxycycline daily from E0-E14 only, collected at NB stage. J, J’, J” and N, N’, N”. RBPj^ind^ embryos injected with doxycycline daily from E14-NB only, collected at NB stage. All data in this Figure are quantified in Figure 8. Scale bars: 100 μm

Having normal NOTCH signaling until E14 then withdrawing doxycycline resulted in a partial rescue of medullary phenotypes at both E16.5 and NB stages (Figs. 7 and 8). At E16, medullary area as measured by UEA-1+ cells was normal (Figs. 7F’, 8A), although UEA-1 intensity had started to decline (Fig. 8B), and both the number and intensity of CLD3,4+ cells was also less than controls (Figs. 7F, 8C,D). PLET-1 staining was also similar to controls (Figs. 7A’, C’; 8E). Thus, just 2 days after withdrawing NOTCH responsiveness mTEC markers had begun to decline. By the NB stage, UEA-1+ area and PLET1 intensity had begun to decline, and UEA-1 intensity remained similar to E16.5 (Fig. 7I, I’, M’); these phenotypes were all improved relative to uninduced RBPj mutants, but remained less than controls (Fig. 8A, B, E). CLD3,4 staining remained similar to that seen at E16.5, and now were also similar to RBPj mutants, in which CLD3,4+ ‘escapers’ have started to accumulate (Figs. 7M; 8C, D). Thus, NOTCH signaling prior to E14 appears to be sufficient to establish an mTEC pool, but it fails to either expand or be maintained properly after doxycycline withdrawal and removal of NOTCH signaling.

**Figure 8.**
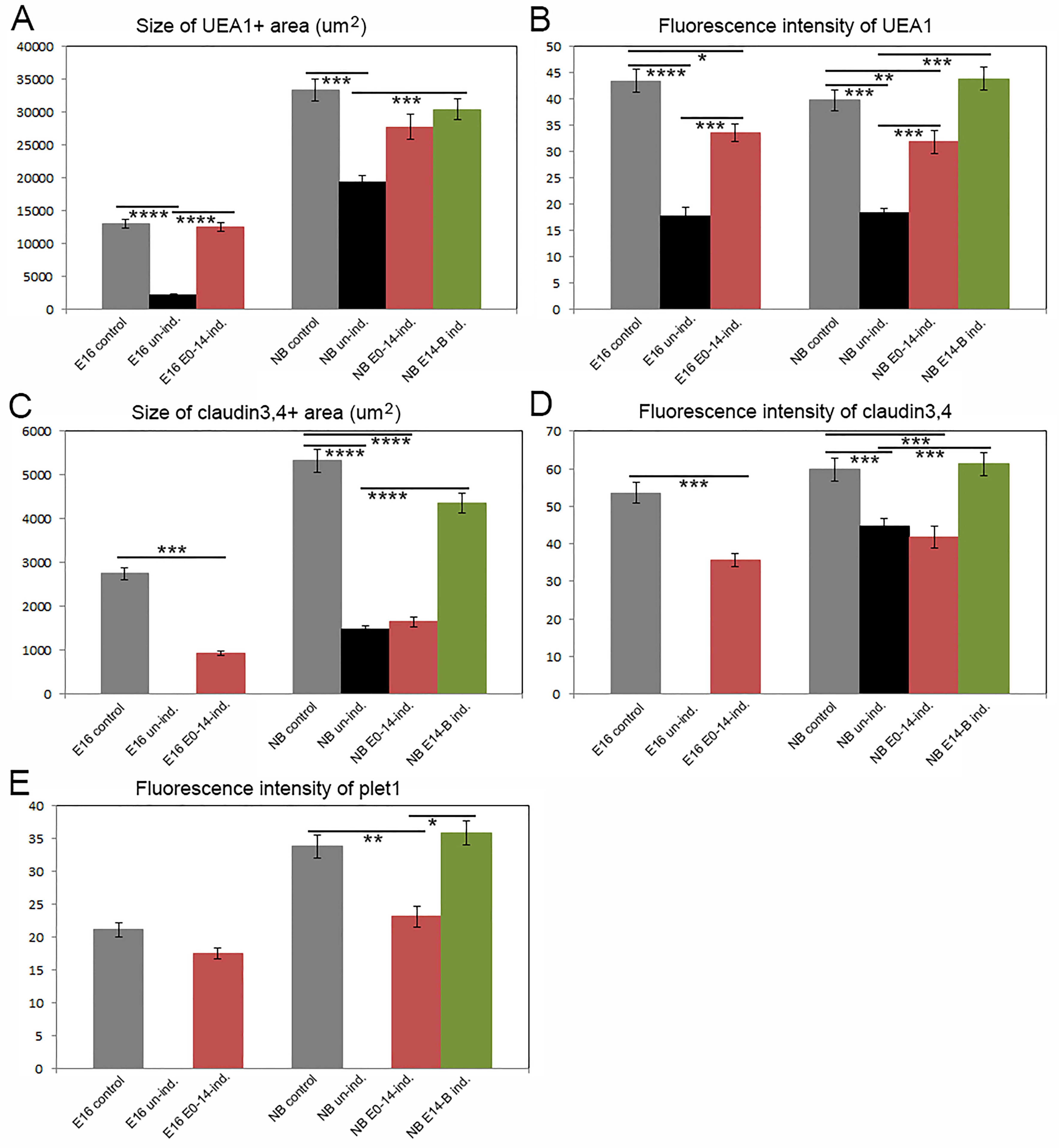
Quantification of TEC marker expression in temporal requirement experiments (see Figure 7). (A,B) Size and fluorescence intensity of UEA1+ area. (C,D) Size and fluorescence intensity of CLD3,4+ area. (E) Fluorescence intensity of PLET1+ cells. All quantification was performed using ImageJ (NIH). \*\*\**P* ≤ 0.0001, \*\*\**P* ≤ 0.001, \*\**P* ≤ 0.005, \**P* ≤ 0.01. n > 3.

In contrast, restoration of NOTCH signaling responsiveness beginning at E14 and continuing until birth substantially restored medullary phenotypes at the NB stage. UEA-1, CLD3,4, and PLET1 intensity were all similar to controls, and significantly increased relative to both uninjected and E0-14 injected samples (Figs. 7J, J’, M, M’; 8A, B, D, E). Only the number of CLD3,4+ cells (measured as area) remained below controls, although was significantly improved relative to uninduced and E0-14 injected samples (Fig. 8C). Furthermore, in both E0-14 and E14-NB samples, CLD3,4 and PLET-1 staining was largely non-overlapping, similar to controls (Fig. S4), and distinct from the maintenance of overlapping staining seen in E16.5 *Foxn1*^*Cre*^*;Notch1*^*fx/fx*^ mutants (Fig. 2), demonstrating that progression from PLET-1+CLD3,4+ to expressing only one or the other marker is NOTCH1-dependent.

These data suggest that NOTCH signaling is required not only for initial mTEC lineage specification, but also for maintenance and/or expansion of the mTEC progenitors throughout fetal stages. These data are also consistent with the possibility that mTEC progenitors can be continue to be specified at later fetal stages.

### Lineage analysis of active Notch signaling in the fetal thymus

We used two NOTCH1 activity-trap mouse lines to trace the lineage of TECs experiencing relatively high (N1IP::Cre^LO^) or lower (N1IP::Cre^HI^) levels of NOTCH1 activation^28^. In these two strains, the NOTCH1 intracellular domain was replaced with Cre, such that NOTCH1 signaling triggers proteolytic cleavage and Cre is able to move to the nucleus. We used these two strains to activate a CAG-tdTomato reporter^29^ to permanently label cells receiving a NOTCH1 signal and their progeny. Co-staining the resulting fetal thymi with TEC markers allowed us to identify all TECs that arise from N1IP::Cre;tdTomato^+^ cells through ontogeny. Interestingly, we observed different patterns of NOTCH1 signaling lineage history in the fetal thymus using these two lineage reporter lines (Figs. 9 and 10).

**Figure 9.**
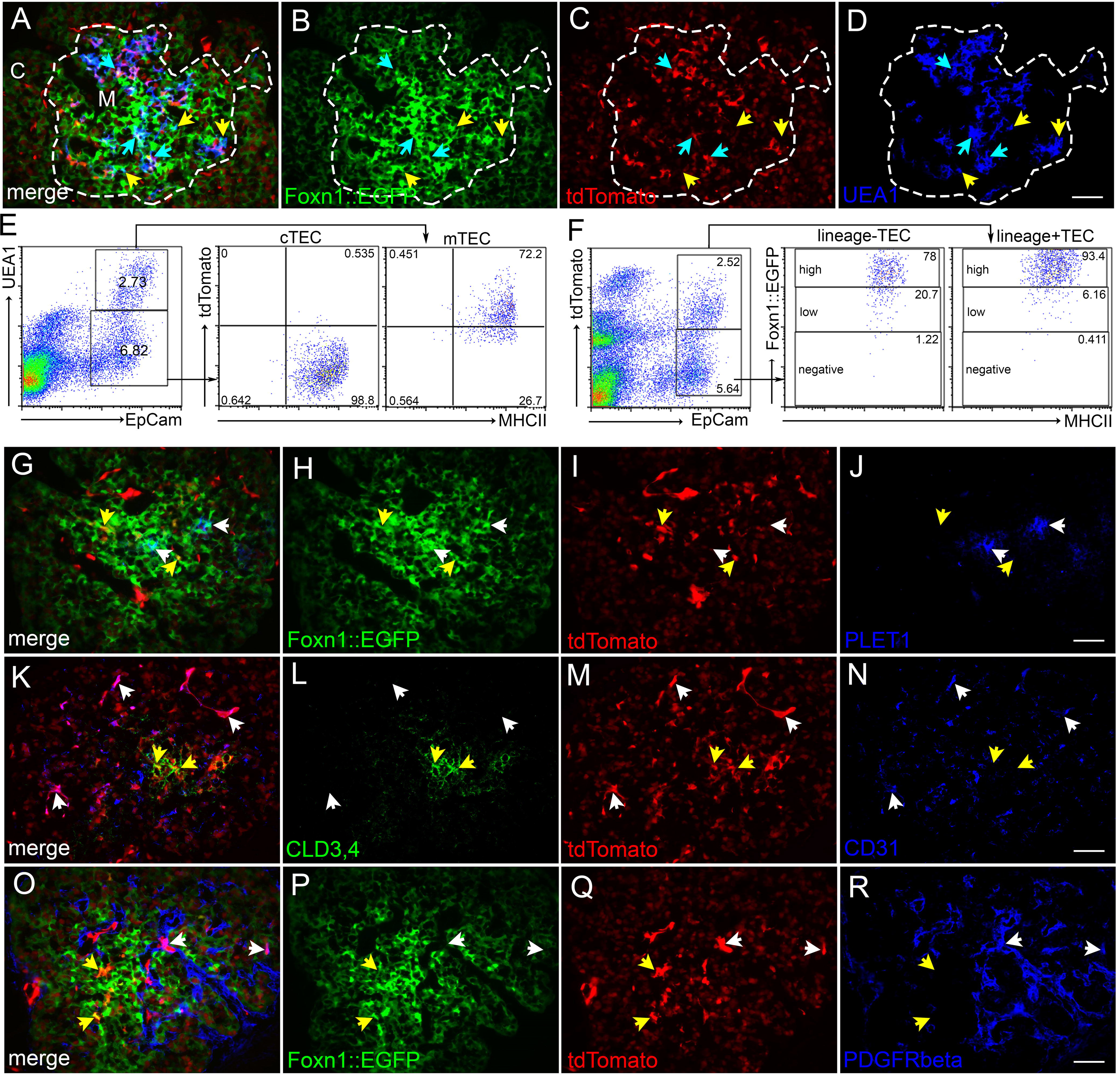
Notch1 signaling lineage tracing in TEPCs: N1IP::Cre^LO^;tdTomato. (A-D) Immunofluorescence of E14.5 N1IP::Cre^LO^;tdTomato;Foxn1::EGFP thymus for expression of Foxn1::EGFP (green; B), tdTomato (red; C) and UEA1 (blue; D). Dashed line outlines medulla. Cyan arrows, GFP^+^;tdTomato^+^;UEA1^+^ cells; yellow arrows, GFP^+^;tdTomato^-^;UEA1^+^ cells. (E) Flow cytometric analysis of newborn N1IP::Cre^LO^;tdTomato thymus stained for EpCam, UEA1 and MHCII, showing percentage of UEA1^+^;MHCII^hi^ mTECs and UEA1^-^;MHCII^hi^ cTECs that express the N1IP::Cre^LO^;tdTomato reporter. (F) Flow cytometric analysis of newborn N1IP::Cre^LO^;tdTomato;Foxn1::EGFP thymus stained for EpCam and MHCII showing Foxn1::EGFP levels in the EpCam^+^;N1IP::Cre^LO^;tdTomato^+^ and EpCam^+^;N1IP::Cre^LO^;tdTomato^-^ TEC populations. (G-J) Immunofluorescence of E14.5 N1IP::Cre^LO^;tdTomato;Foxn1::EGFP thymus for Foxn1::EGFP (green; H), tdTomato (red; I) and Plet1 (blue; J). White arrows, GFP^+^;tdTomato^-^;PLET1^+^ TEPCs; yellow arrows, GFP^+^;tdTomato^+^;PLET1^-^ TECs. (K-N) Immunofluorescence of E14.5 N1IP::Cre^LO^;tdTomato thymus for CLD3,4 (green; L), tdTomato (red; M) and CD31 (blue; N). White arrows, CLD3,4^-^;tdTomato^-^;CD31^+^ endothelial cells; yellow arrows, CLD3,4^+^;tdTomato^+^;CD31^-^ mTEPCs. (O-R) Immunofluorescence of E14.5 N1IP::Cre^LO^;tdTomato;Foxn1::EGFP thymus for Foxn1::EGFP (green; P), tdTomato (red; Q) and PDGFR-β (blue; R). White arrows, GFP^-^;tdTomato^+^;PDGFR-β^+^ pericytes; yellow arrows, GFP^+^;tdTomato^+^;PDGFR-β^-^ TECs. Scale bars, 50 μm. C, cortex. M, medulla. n > 3 for IHC; n > 5 for flow cytometry.

**Figure 10.**
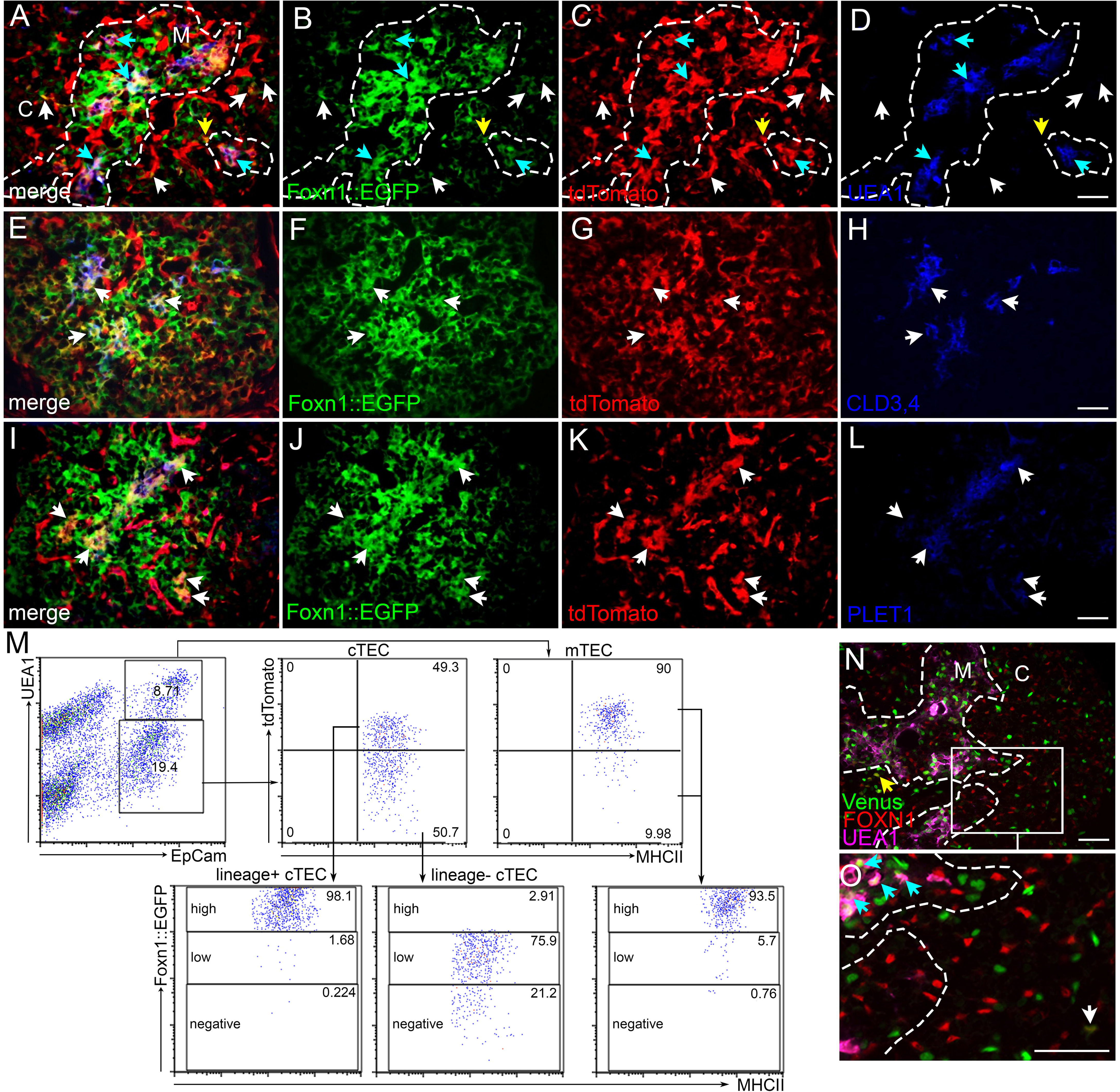
Notch1 signaling lineage tracing in TEPCs: N1IP::Cre^HI^;tdTomato. (A-D) Immunofluorescence of E14.5 N1IP::Cre^HI^;tdTomato;Foxn1::EGFP thymus for expression of Foxn1::EGFP (green; B), tdTomato (red; C) and UEA1 (blue; D). White arrows, GFP^+^;tdTomato^+^;UEA1^-^ cTECs; yellow arrow, GFP^+^;tdTomato^+^;UEA1^-^ cell at the cortico-medullary junction; cyan arrows, GFP^+^;tdTomato^+^;UEA1^+^ mTECs. Dashed line outlines medulla. (E-H) Immunofluorescence of E14.5 N1IP::Cre^HI^;tdTomato;Foxn1::EGFP thymus for, Foxn1::EGFP (green; I), tdTomato (red; J) and Cld3,4 (blue; K). Arrows, GFP^+^;tdTomato^+^;CLD3,4^+^ cells. (I-L) Immunofluorescence of E14.5 N1IP::Cre^HI^;tdTomato;Foxn1::EGFP thymus for, Foxn1::EGFP (green; M), tdTomato (red; N) and PLET1 (blue; O). Arrows, GFP^+^;tdTomato^+^;PLET1^+^ cells. (M) Flow cytometric analysis of newborn N1IP::Cre^HI^;tdTomato;Foxn1::EGFP thymus stained for EpCam, UEA1 and MHCII showing percentage of UEA1^+^ mTECs and UEA1^-^ cTECs that express the N1IP::Cre^HI^;tdTomato reporter. Lower plots show Foxn1::EGFP expression levels in UEA1^-^;tdTomato^+^ cTECs and UEA1^-^;tdTomato^-^ cTECs. (N,O) Immunofluorescence of E16.5 CBF:H2B-Venus thymus for, FOXN1 (red) and UEA1 (magenta). Yellow arrow, Venus+; FOXN1+; UEA1-TEC at the cortico-medullary junction; white arrow, Venus+; FOXN1+; UEA1-TEC in the cortex; cyan arrows, Venus+; FOXN1+; UEA1+ mTECs. Box in (N) is zoomed area in (O). Dashed line outlines medulla. Scale bars, 50μm. C, cortex. M, medulla. n > 3 for IHC; n > 5 for flow cytometry.

Analysis of the N1IP::Cre^LO^;tdTomato reporter (Fig. 9) at E14.5 identified only those cells that either themselves or their progenitors had experienced a *high* level of NOTCH1 signaling prior to or at that stage. To assess TEC positive for this marker, we used both the tdTomato reporter and Foxn1::GFP to identify TEC (N1IP::Cre^LO^;tdTomato;Foxn1::EGFP) (see Fig. S3 for gating controls used for these two markers). At E14.5, a subset of medullary TECs marked by UEA1 staining were lineage-positive, (blue arrows, Fig. 9A-D), although a substantial fraction of mTEC were lineage-negative (yellow arrows, Fig. 9A-D). Consistent with this result, flow cytometry showed that around 75% of MHCII^hi^;UEA1^+^ mTECs expressed the N1IP::Cre^LO^;tdTomato reporter at the newborn stage (Fig. 9E, right panel), while fewer than 1% of MHCII^hi^;UEA1^-^ cTECs had experienced high levels of NOTCH1 activity (Fig. 9E, middle panel). Almost all lineage-positive TECs (N1IP::Cre^LO^tdTomato^+^EpCAM^+^) and lineage-negative TECs (N1IP::Cre^LO^;tdTomato^-^) were Foxn1::EGFP^+^MHCII^+^ (Fig. 9F), confirming the TEC identity of the cells. In terms of progenitors, CLD3,4^+^ cells expressed the N1IP::Cre^LO^;tdTomato reporter (yellow arrows, Fig. 9L,M), whereas PLET1^+^ cells did not (white arrows in Fig 9H-J). These data are consistent with our CBF:H2B-Venus reporter data (Fig. 1K,L) showing that the mTEPC pool is undergoing active NOTCH signaling; these data specifically show that CLD3,4^+^ cells have experienced a high level of NOTCH1 signal. Lineage-positive non-TEC cells (N1IP::Cre^LO^tdTomato^+^ cells negative for TEC markers) were vascular-associated, as indicated by co-expression with CD31 (white arrows, Fig. 9K-N) and PDGFR-β (white arrows, Fig. 9O-R).

Next, we assessed the expression pattern of the N1IP::Cre^HI^;tdTomato reporter in the thymus at E14.5, which reports a broader range of NOTCH1 signaling (Fig. 10). Almost all UEA1^+^ mTECs expressed the N1IP::Cre^HI^;tdTomato reporter at E14.5 by IHC (cyan arrows in Fig. 10C,D) and at the newborn stage by flow cytometric analysis (Fig. 10M). Consistent with our other expression, signaling, and lineage results, all CLD3,4^+^ cells were N1IP::Cre^HI^;tdTomato^+^ (arrows in Fig. 10E-H). However, in contrast to the results from the N1IP::Cre^LO^ reporter, most or all PLET1^+^ cells were also positive for this reporter (arrows in Fig. 10I-L). These results support a model in which all TEPCs have experienced at least low levels of NOTCH1 signaling, while those receiving a high level of signaling commit to the mTEC fate.

Analysis of this reporter in cTECs showed that some lineage-positive Foxn1::GFP^+^ TECs could also be detected in the cortex (white arrows in Fig. 10A-D). Flow cytometry revealed that around half of all cTECs (EpCam^+^UEA1^-^) were tdTomato^+^ at the newborn stage (Fig. 10M). This finding reveals a previously unidentified split in the cTEC population, based on history of NOTCH1 signaling. Essentially all (> 98%) of the NOTCH1 lineage-positive cTECs (EpCam^+^UEA1^-^N1IP::Cre^HI^tdTomato^+^) were Foxn1::EGFP^hi^ (Fig. 10M). However, none of the NOTCH1 lineage-negative cTECs expressed a high level of Foxn1::EGFP (Fig. 10M). These Foxn1::EGFP low cells also had lower MHCII surface levels than the Foxn1::EGFP high cells (MFI 256, SD = 18.73 vs. MFI 360, SD = 31.53; *P* = 0.008). Thus, the expression levels of FOXN1 and MHCII are correlated in these cell populations consistent with previous studies, and the lower levels are also consistent with a less differentiated phenotype.

Finally, to assess the level of *current or recent* as opposed to a *history* of NOTCH signaling, we analyzed CBF:H2B-Venus expression at E16.5. While a substantial fraction of mTECs and all CLD3,4^+^ mTECs were Venus^+^FOXN1^+^, there were only rare Venus^+^FOXN1^+^ cells in the cortex (Fig. 10N,O). This result suggests the existence of two distinct populations of cells within the lineage-negative cTECs, and suggests that the NOTCH lineage-positive cTECs may arise from a relatively small population of cTECs undergoing active NOTCH signaling.

In summary, we have generated a fate map of NOTCH1 signaling during TEC ontogeny using two NOTCH1 activity-trap mouse lines. Our data reveal that all mTECs, but only a subset of cTECs, have experienced NOTCH1 signaling during fetal thymus development.

## Discussion

Thymic epithelial cells (TECs) represent the major functional component of the thymus, yet the mechanisms controlling their differentiation during fetal development remain largely unknown, particularly in terms of lineage specification and progenitor cell maintenance. In the current study, we provide evidence that NOTCH1 signaling is required to specify the lineage-restricted mTEC progenitor pool in the fetal thymus. We show that all mTEPCs in the fetal thymus exhibit active NOTCH1 signaling from early in organogenesis, and have a lineage history of high levels of NOTCH signaling. Ablation of *Notch1* in TECs results in fewer TEPCs and causes a block in specification of mTEC progenitors, as *Notch1* null TEC are unable to contribute to the mTEC lineage after mosaic deletion. In contrast, NOTCH1 activation in TECs results in an expansion of the TEPC pool, but then subsequent mTEC differentiation is also blocked. These data indicate that NOTCH signaling is required for specification of mTEC progenitors, and promotes their expansion, but that NOTCH signaling must cease for mTEC differentiation to mature phenotypes to occur. The fact the removal of NOTCH signaling in TEC after E14 results in progressive loss of the mTEC population also suggests that NOTCH signaling is required for maintenance of mTEC progenitors, or for their proliferation. The similarity in phenotypes from targeting RBPj and NOTCH1 suggests that at fetal stages *Notch1* is the major mediator of NOTCH signaling in TEC. A parallel study in the Blackburn lab targeting RBPj and thus globally affecting NOTCH signaling came to a similar conclusion (Liu, et al., companion paper).

The developmental origins of separate cortical and medullary TEC lineages and the existence and identity of bipotent TEC progenitors remains controversial. Whether they arise from a common bipotent or individual lineage-restricted progenitors is still uncertain, with evidence for both^11-16^. Furthermore, it is still unclear exactly when and how the fetal and adult TEC progenitor populations arise and what their relationships may be. Our data do not definitively prove either the bipotent or the individual lineage-restricted progenitor model, but do provide clear indications of how different lineages are related, and show that mTEC and cTEC require different signals for specification.

We identify NOTCH1 as a key molecule required for the establishment and expansion of the mTEC progenitor pool. Our functional studies revealed that NOTCH1 pathway inhibition or activation both affected the mTEPC pool in the fetal thymus. Our data are consistent with a model in which NOTCH1 signaling acts on an early fetal bipotent progenitor that is PLET1^+^CLD3,4^+^, which gives rise to a PLET1^-^CLD3,4^+^ mTEC-specific TEPC pool that has experienced high levels of NOTCH signaling, sometime between E13.5 and E16.5. Whether this PLET1^+^CLD3,4^+^ TEPC also gives rise to the cortical lineage is not clear; as other lineage studies have suggested that all TECs arise from a progenitor expressing cortical markers^13,14^. However, it is clear that cTECs do not all experience NOTCH signaling, at least not at levels we can detect with our lineage reporters, and that cTEC in general can develop in the absence of NOTCH signaling. In either case, our data indicate that a bipotent progenitor would likely itself not experience NOTCH signaling, although its immediate daughter cells could.

We propose a model in which NOTCH1 signaling is required to generate the mTEPC pool during fetal thymus development (Fig. 11). Lineage restriction of these cells occurs according to whether or not the bipotent progenitor itself, or its daughter cells, experience high levels of NOTCH1 signaling. In this model, all TECs arise from a common bipotent progenitor cell, although it is also formally possible that the PLET1^+^CLD3,4^+^ TEPC population contains separate cortical and medullary progenitors. Regardless, those cells that do receive a NOTCH1 signal will become PLET1^-^CLD3,4^+^ mTEC lineage-restricted TEPCs; those that do not become cTEC, either by default or under the influence of a second unknown signal. Thus, when *Notch1* is deleted from TECs (as in the *Foxn1*^*Cre*^*;Notch1*^*fx/fx*^ and *Foxg1*^*Cre*^*;Notch1*^*fx/fx*^ models presented here) the PLET1^+^CLD3,4^+^ TEPCs fail to down regulate *Plet1* and the mTEPC lineage-restricted pool is not generated. Our data also show that *Notch1* must be down regulated for differentiation of the PLET1^-^CLD3,4^+^ cells into more mature mTECs, consistent with previous reports^7^. Thus, in our *Foxn1*^*Cre*^*;Rosa*^*N1-IC*^ over-expression model, prolonged NOTCH1 signaling prevents mTEC differentiation and fewer mature AIRE^+^ mTECs are made.

**Figure 11.**
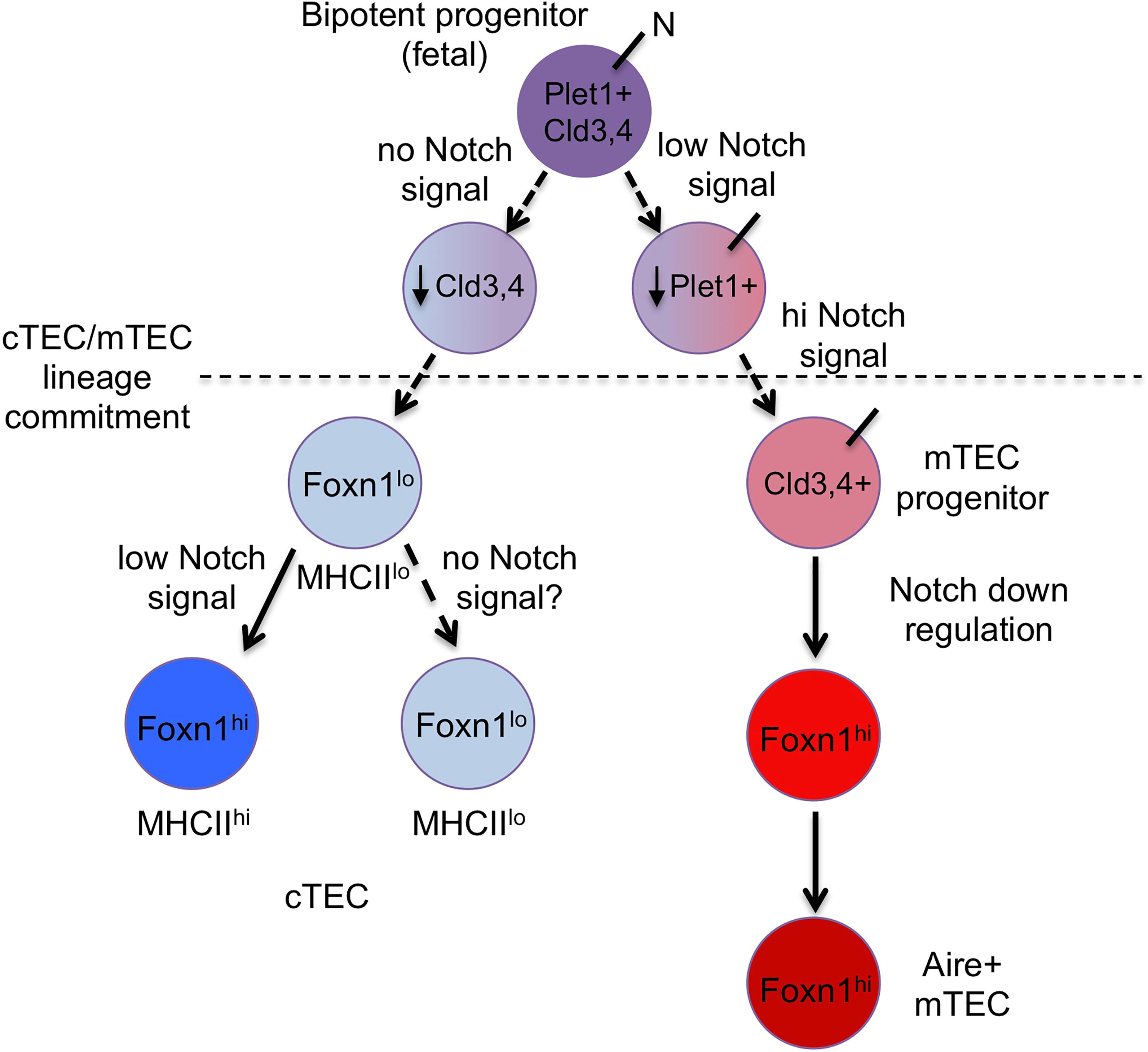
Model for the role of Notch1 signaling during fetal TEC development. In this model, all fetal TECs derive from a common PLET1^+^;CLD3,4^+^ progenitor pool that will then become lineage-restricted into either mTEPCs or cTEPCs. While the bipotent progenitor itself does not experience NOTCH signaling, immediate progeny that experience low levels of NOTCH signaling down regulate PLET-1 and up regulate CLD3,4, committing to the mTEC lineage; these mTEPCs then experience high levels of NOTCH signaling to drive initial expansion and differentiation. *Notch1* expression must then be down regulated in those cells for mTEC differentiation to proceed functional AIRE^+^ mTECs. The progeny of PLET1^+^ cells that do not receive a NOTCH1 signal will down regulate CLD3,4 expression and progress to the cTEC lineage. At some point during their differentiation, a separate exposure to low NOTCH signaling results in up-regulation of *Foxn1*, presumably leading to cTEC maturation. It is also possible that the cTEC lineage splits into two different functional populations depending on exposure to low level NOTCH signaling (dotted arrow); in the absence of more cTEC markers and functional information, these two possibilities cannot be distinguished.

Our fate mapping lineage analysis showed that only half of fetal cTECs have experienced NOTCH1 signaling, and that these cTECs have uniformly higher levels of *Foxn1* and MHCII expression than those that are NOTCH lineage-negative. These data indicate that NOTCH signaling may also play a role in cTEC differentiation that is distinct from the mTEC role, uncovering a previously unidentified diversity within cTEC based on having experienced NOTCH signaling (Fig. 9). Compared to mTECs, little is known about the cTEC lineage and its development during ontogeny. As these two lineage-negative and lineage-positive populations also differ in their levels of *Foxn1* and MHCII expression, it is reasonable to conclude that these populations may be distinct either in their level of maturity or their function. Although we did not detect an obvious change in cTECs in our *Notch1* deletion model, the relative lack of cTEC markers means that we have little power to do so based on known markers. As a result, we can only speculate at this point what the relationship between these two cTEC subsets may be. As the lineage positive cTECs cannot give rise to lineage negative cTECs due to the nature of our reporters, either the lineage-negative cTECs must give rise to lineage-positive cTECs upon experiencing NOTCH signaling, or the two populations have to arise independently. Regardless, this result indicates that low level NOTCH signaling acts on cTECs, and opens new avenues of investigation into cTEC differentiation.

NOTCH signaling functions via cell-cell contact, therefore the NOTCH1 signal that TECs experience must be triggered by ligands expressed on adjacent cells. But what are these cells? What cells express the ligand(s), and what are the ligands? The cells could be other TECs, thymocytes, endothelial cells and/or neural crest-derived mesenchymal cells. It has been suggested that thymocytes are at least one source of ligand, and that an interaction between these two cell types is required for TEC development^6^. In the current study, we first observed active NOTCH1 signaling in Foxn1^+^ cells at early E11.5, which is coincident with the first wave of lymphocyte entry to the primordium^30^, although it is clear in our data that TECs are not adjacent to thymocytes when undergoing NOTCH signaling. Of note, at this early stage there are few cellular sources of NOTCH ligands, and the most likely source based on our expression data are other fetal TECs, which express multiple NOTCH ligands, including *Jagged1* and *Delta-like4*^5,6,30,31^ (Liu, et al, co-submitted paper). Whether the specific ligands and their cellular source change during ontogeny, or have functional consequences for TEC biology, remain to be determined.

## Methods

### Mice

#### At UGA

*Notch1*^*flox*^ (Stock No. 006951), *Rosa*^*N1-IC*^ (Stock No. 008159), *CBF:H2B-Venus* (Stock No. 020942) and *CAG-tdTomato* (Stock No. 007909) mice were obtained from The Jackson Laboratories (Bar Harbor, ME). N1IP::Cre^HI^ and N1IP::Cre^LO^ strains were a gift from Dr. Raphael Kopan (Cincinnati Children’s Hospital Medical Center, Cincinnati, OH)^28^. Foxn1::EGFP (enhanced green fluorescent protein) mice were a gift from Dr. Thomas Boehm (Max Planck Institute of Immunobiology, Freiburg, Germany)^32^. *Foxn1*^*Cre*^ and *Foxa2* ^*Cre*^ strains have been described elsewhere^20,33^. All colonies were maintained on a majority C57BL6/J genetic background. Noon on the day of detecting a vaginal plug was designated embryonic day 0.5 (E0.5), and confirmed by morphological features.

All mice and embryos were genotyped by PCR using DNA extracted from tail tissue. EGFP primer sequences were: fwd, GTT CAT CTG CAC CAC CGG C; rev, TTG TGC CCC AGG ATG TTG C. Primer sequences for *Notch1*^*flox*^, *Rosa*^*N1-IC*^, CBF:H2B-Venus, CAG-tdTomato, *Foxn1*^*Cre*^ (*Foxn1*^*ex9cre*^, Stock No. 018448), and *Foxg1*^*Cre*^ (Stock No. 006084) strains are available from The Jackson Laboratories (Bar Harbor, ME). In all cases, Cre negative animals or embryos were used as littermate controls. n-values for all experiments are shown in figure legends.

All experiments involving animals were performed with approval from the UGA Institutional Animal Care and Use Committee.

#### At Toronto

RBPj-inducible (RBPj^ind^ or RBPj^fx/fx^;Rosa^rtTA^;Tet^on^-RBPj-HA) mice, described elsewhere ^27^, were bred to FoxN1^cre^ mice (RBPj^fx/fx^;Foxn1^Cre^;Rosa^rtTA^;Tet^on^-RBPj-HA) and maintained in the Comparative Research Facility of the Sunnybrook Research Institute under specific pathogen-free conditions. All animal procedures were approved by the Sunnybrook Research Institute Animal Care Committee and performed in accordance with the committee’s ethical standards. For induction of Notch responsiveness, pregnant mice were given 1 mg/ml Doxycycline (Sigma-Aldrich) in drinking water supplemented with 5% Splenda *ad libitum*.

### Immunofluorescence and histology

For cryosectioning, mouse embryos were snap frozen in liquid nitrogen and stored at −80°C. Tissues were sectioned at 8 μm and fixed in ice-cold acetone for 2□min. Tissues were rinsed with phosphate buffered saline (PBS), blocked with 10% donkey serum in PBS for 30□min at room temperature, then incubated with appropriate primary antibodies overnight at 4°C: anti-cleaved NOTCH1 (Cell Signaling Technologies, 4147, 1:200), anti-NOTCH1 (Origene, EP1238Y, 1:200), anti-Foxn1 (Santa Cruz, G-20, 1:200), anti-CD31 (BD, MEC13.3, 1:100), anti-PDGFR-β (R&D Systems, AF1042, 1:50), anti-Ikaros (Santa Cruz, M-20, 1:200), anti-GFP (Abcam, ab13970, 1:200), anti-Plet1 (rat supernatant from cell line ID4-20), anti-Claudin3 (Life Technologies, 34-1700, 1:200), anti-Claudin 4 (Life Technologies, 36-4800, 1:200), anti-β5t (MBL, PD021, 1:200), anti-CD205 (BioLegend, 138202, 1:200), anti-Aire (Santa Cruz, M-300, 1:200), anti-K5 (Covance, AF138, 1:1,000), anti-K8 (rat supernatant, Troma1), anti-K14 (Covance, AF64, 1:1,000) or UEA1 lectin (Vector Labs, X0922, 1:400). Secondary detection was performed with donkey anti-primary species. For *N1IP::Cre;tdTomato;Foxn1::EGFP* observation, tissues were fixed in 4% paraformaldehyde (PFA) in PBS for 5□min at 4°C, washed with PBS followed by 10% sucrose/PBS for 1□h, then 30% sucrose/PBS overnight. Tissues were embedded in OCT and stored at −80°C until sectioning. Sections were examined by fluorescent microscopy using a Zeiss Axioplan 2 microscope (Thornwood, NY).

For paraffin sectioning, tissues were collected and fixed in 4% PFA for 2-3 h. Tissues were dehydrated through an ethanol series (70%, 80%, 90%, 96%, 100%) and embedded in paraffin wax using standard procedures. Sections (8□μm) were cut and rinsed in xylene before rehydration through a reverse ethanol series. Antigen retrieval was performed by boiling slides in 10 mM sodium citrate buffer, pH 6, for 30 min. Sections were stained using appropriate primary and secondary antibodies as described above, and imaged using fluorescence microscopy.

Hematoxylin and eosin (H&E) staining was performed on paraffin sections using standard procedures, then imaged on a Zeiss Axioplan microscope (Thornwood, NY).

### Flow cytometry

For TEC analysis, fetal or newborn stage thymi were dissected and digested in 1 mg/mL collagenase/dispase (Roche, Basel, Switzerland), and passed through a 100-μm mesh to remove debris. Thymi were processed individually, before genotyping. PE-Cy7 conjugated anti-CD45 (BioLegend, 30-F11, 1:150) and APC-conjugated anti-EpCam (BioLegend, G8.8, 1:150) were used to isolate TEC populations. UEA1 lectin, anti-Claudin 3 and anti-MHCII (M5/114.15.2, BioLegend, 1:150) were used in the TEC analysis. Cells were refixed in 1% PFA/PBS and analyzed using a CyAn ADP Flow Cytometer (Beckman Coulter, Miami, FL). Data were collected using a four-decade log amplifier and stored in list mode for subsequent analysis using FlowJo Software (Tree Star, Ashland, OR).

Thymocytes were harvested from individual fetal or newborn stage thymi and suspended in FACS buffer (PBS with 2% fetal bovine serum (FBS)). Thymi were processed individually, before genotyping. Cells were incubated with conjugated monoclonal antibodies CD4-FITC (BioLegend, GK1.5, 1:150), CD8-PE (BioLegend, 53-6.7, 1:150), CD25-APC (BD, PC61, 1:150) or CD44-PerCP (BioLegend, IM-7, 1:150), at 4°C for 30□min, washed, and fixed with 1% PFA (EM Sciences, Ft. Washington, PA) before analysis on a CyAn ADP Flow Cytometer (Beckman Coulter, Miami, FL). Data were collected on using a four-decade log amplifier and were stored in list mode for subsequent analysis using FlowJo Software.

### Cell isolation and genomic PCR

E15.5 thymi were harvested and processed individually to generate a single cell suspension (as described above). TEC populations were isolated based on staining with PE-Cy7 conjugated anti-CD45, APC-conjugated anti-EpCam, UEA1 lectin and anti-MHCII as described in the text. DNA was purified from sorted cell populations using QIAamp DNA Mini kit (QIAGEN). PCR was performed using the following primer sequences: fwd-1 (undeleted allele), TAC TTA GAG CGG GGC AGA GA; fwd-2 (deleted allele), CTG AGG CCT AGA GCC TTG AA; rev (both deleted and undeleted alleles), ACT CCG ACA CCC AAT ACC TG.

### Statistics

Data are presented as mean ± S.D. N values were at least 3 for each genotype in each experiment and are indicated in the text and/or Figure legends. Comparisons between two groups were made using Student’s *t* test, multiple comparisons used ANOVA. *P* < 0.05 was considered significant.

## Acknowledgements

We thank E. Richie for providing the K8 and PLET1 antibodies, and C.C. Blackburn and E. Richie for reading the manuscript and helpful discussions. We thank J. Nelson in the Center for Tropical and Emerging Global Diseases Flow Cytometry Facility at the University of Georgia for flow cytometry and cell sorting technical support. This study was supported by grant # R21 AI107465 to NM from the National Institutes of Health, and a Canadian Institutes for Health Research (CIHR) grant (FND-154332) to JCZP. ELYC was supported by an Ontario Graduate Scholarship, and JCZP is supported by a Canada Research Chair in Developmental Immunology.

## Author contributions

NM and JCZF designed the experiments; JL, JG, ELYC and LW performed the experiments and generated the data. All authors participated in data analysis. JG, JL and NM prepared the manuscript.

## Competing financial interests

The authors declare no competing financial interests.

**Figure S1. Fewer TEPCs in the *Foxg1***^***Cre***^***;Notch1***^***fx/fx***^ **fetal thymus.** Immunofluorescence of E14.5 (A,B) and E18.5 (C,D) *Foxg1*^*Cre*^*;Notch1*^*fx/fx*^ mutant (B,D) and control (A,C) thymus for expression of CLD3,4 (red), PLET1 (green) and UEA1 (blue). Scale bars, 50 μm. n > 3.

**Figure S2. TEC organization and differentiation are affected in the *Foxg1***^***Cre***^***;Notch1***^***fx/fx***^ **fetal thymus.** (A,B) Immunofluorescence of E12.5 *Foxg1*^*Cre*^*;Notch1*^*fx/fx*^ mutant (B) and control (A) thymus for expression of K5 (red), K8 (green) and UEA1 (blue). (C,D) Immunofluorescence of E18.5 *Foxg1*^*Cre*^*;Notch1*^*fx/fx*^ mutant (D) and control (C) thymus for expression of K5 (red), K8 (green). (E,F) Immunofluorescence of E18.5 *Foxg1*^*Cre*^*;Notch1*^*fx/fx*^ mutant (F) and control (E) thymus for expression of AIRE. (G) Flow cytometric analysis of intrathymic thymocytes isolated from E18.5 *Foxg1*^*Cre*^*;Notch1*^*fx/fx*^ mutant and control thymi stained for CD4, CD8, CD44 and CD25 subsets. Scale bars, 50 μm. n > 3 for IHC; n > 5 for flow cytometry.

**Figure S3. Gating controls for flow cytometric analysis of thymic cells isolated from newborn N1IP::Cre**^**LO**^**;tdTomato;Foxn1::EGFP and N1IP::Cre**^**HI**^**;tdTomato;Foxn1::EGFP mice.** Cells were divided into FOXN1 high, low, and very low/negative for the analyses shown in Figures 6 and 7 based on these gates.

**Figure S4. Restoring NOTCH signaling receptivity in TECs rescues mTEPC generation.** (A-D) Immunofluorescence of E16 (A-C) or NB (D-G) thymi collected from controls RBPj^fx/+^;Foxn1^Cre^;Rosa^rtTA^;Tet^on^-RBPj-HA (A, D), uninduced RBPj^fx/fx^;Foxn1^Cre^;Rosa^rtTA^;Tet^on^-RBPj-HA (RBPJ^ind^) (B, E), or RBPJ^ind^ mice injected with doxycycline from E0-14 (C, F) or from E14-NB (G), as in Figure 7. Thymi are stained for expression of CLD3,4 (red) and PLET1 (green). White arrows indicate PLET1^+^;CLD3,4^-^ cells; cyan arrows indicate PLET1^-^;CLD3,4^+^ cells; yellow arrows indicate PLET1^+^;CLD3,4^+^ cells. Compare with Figure 2. Scale bars: 50 μm.

